# Modeling implicates inhibitory network bistability as an underpinning of seizure initiation

**DOI:** 10.1101/613794

**Authors:** Scott Rich, Homeira Moradi Chameh, Marjan Rafiee, Katie Ferguson, Frances K. Skinner, Taufik A. Valiante

## Abstract

A plethora of recent experimental literature implicates the abrupt, synchronous activation of GABAergic interneurons in driving the sudden change in brain activity that heralds seizure initiation. However, the mechanisms predisposing an inhibitory network toward sudden coherence specifically during ictogenesis remain unknown. We address this question by comparing simulated inhibitory networks containing control interneurons and networks containing hyper-excitable interneurons modeled to mimic treatment with 4-Aminopyridine (4-AP), an agent commonly used to model seizures *in vivo* and *in vitro*. Our *in silico* study demonstrates that model inhibitory networks with 4-AP interneurons are more prone than their control counterparts to exist in a bistable state in which asynchronously firing networks can abruptly transition into synchrony due to a brief perturbation. We further show that perturbations driving this transition could reasonably arise *in vivo* based on models of background excitatory synaptic activity in the cortex. Thus, these results propose a mechanism by which an inhibitory network can transition from incoherent to coherent dynamics in a fashion that may precipitate seizure as a downstream effect. Moreover, this mechanism specifically explains why inhibitory networks containing hyper-excitable interneurons are more vulnerable to this state change, and how such networks can undergo this transition without a permanent change in the drive to the system. This, in turn, potentially explains such networks’ increased vulnerability to seizure initiated by GABAergic activity.

**Author summary:** For decades, the study of epilepsy has focused on the hypothesis that over-excitation or dis-inhibition of pyramidal neurons underlies the transition from normal brain activity to seizure. However, a variety of recent experimental findings have implicated a sudden synchronous burst of activity amongst inhibitory interneurons in driving this transition. Given the counter-intuitive nature of these findings and the correspondingly novel hypothesis of seizure generation, the articulation of a feasible mechanism of action underlying this dynamic is of paramount importance for this theory’s viability. Here, we use computational techniques, particularly the concept of bistability in the context of dynamical systems, to propose a mechanism for the necessary first step in such a process: the sudden synchronization of a network of inhibitory interneurons. This is the first detailed proposal of a computational mechanism explaining any aspect of this hypothesis of which we are aware. By articulating a mechanism that not only underlies this transition, but does so in a fashion explaining why ictogenic networks might be more prone to this behavior, we provide critical support for this novel hypothesis of seizure generation and potential insight into the larger question of why individuals with epilepsy are particularly vulnerable to seizure.

## Introduction

Epilepsy is a neurological disease distinguished by repeated seizures, often characterized by hyper-excitable and synchronous activity of pyramidal neurons. Epilepsy research is typically divided into studies focused on either seizure initiation [1], propagation [2, 3], or termination [4], as schematized in Fig 1 [5]. Historically, studies of seizure initiation have focused on the hypothesis that hyper-excitability of excitatory cells is the impetus of seizure [5] and associated inhibitory collapse. However, there is as of yet no cellular-based mechanism explaining the transition into seizure [6, 7].

**Fig 1.**
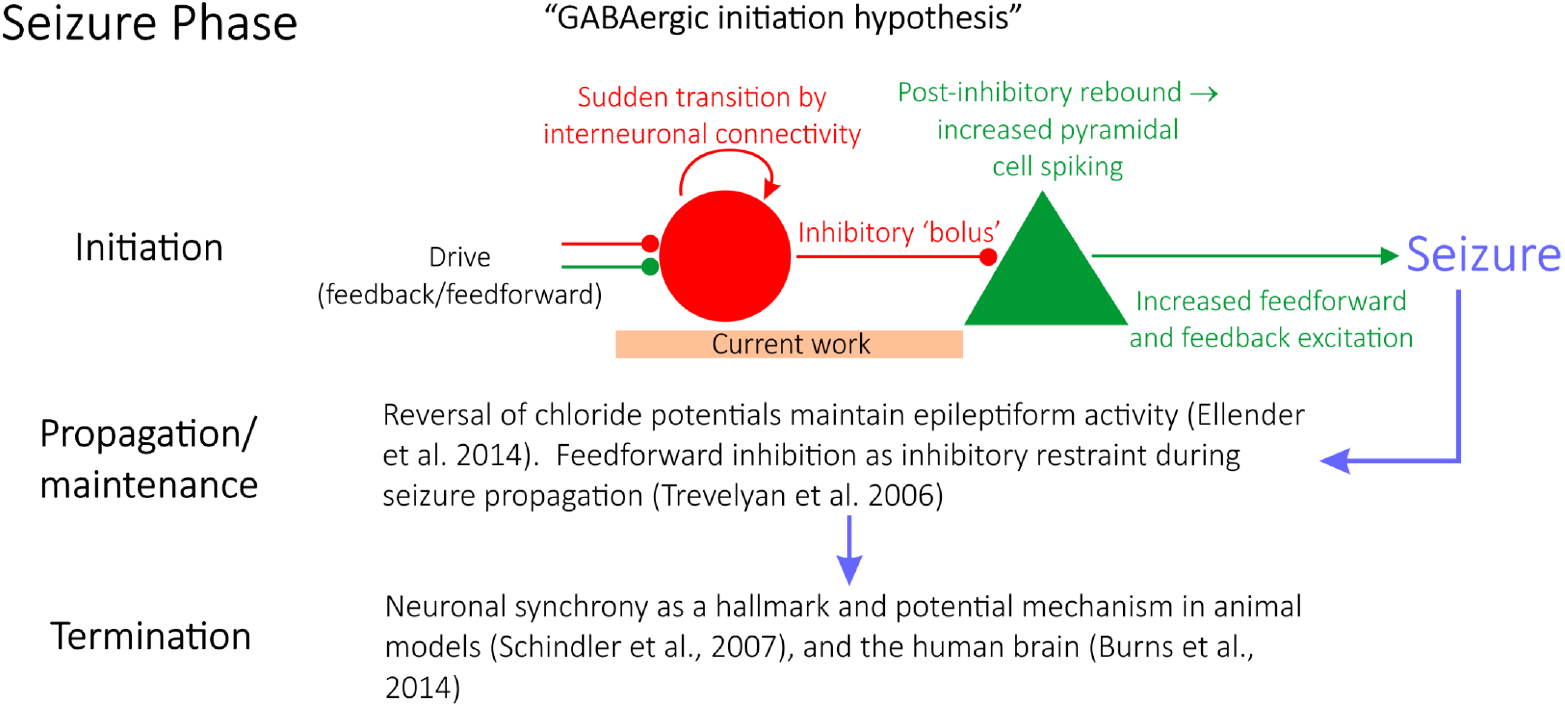
A “GABAergic initiation hypothesis” in the context of the state of epilepsy research. Epilepsy research is divided into studies focusing on seizure initiation, propagation, or termination (illustrated by the rows in this figure and the example studies cited). This paper is interested in seizure initiation in the context of a “GABAergic initiation hypothesis”, schematized on the top row of the figure. The focus of the current work is the sudden transition of interneurons into synchrony, as the articulation of a potential mechanism explaining this transition, alongside a justification as to why networks in a seizure state are more vulnerable to this transition, should be identified in order for this overall hypothesis of seizure initiation to be viable.

Recently, some studies of seizure initiation have shifted focus to over-activity of inhibitory interneurons. This literature has yielded convincing evidence that interneurons serve a causal role in seizure initiation [1, 8–13], laying the groundwork for a novel hypothesis for seizure initiation (a “GABAergic initiation hypothesis”) in which synchronous activation of inhibitory interneurons precipitates the onset of a seizure, as diagrammed in Fig 1 [12]. Given the contemporaneous nature of this hypothesis it is an ideal target for rigorous computational study; here, such research aims to unearth a mechanism explaining the predisposition of inhibitory interneurons in a hyper-excitable environment to suddenly transition into synchrony, the necessary initial step in this hypothesis. We thus focus on the earliest time in the transition to seizure and not aspects of propagation and termination.

The study of inhibitory network synchrony is decades old, dating back to the work of Wang and Rinzel [14]. Various mechanisms have been proposed to explain the generation of oscillations in purely inhibitory networks, the most prominent of which may be the Interneuron Network Gamma (ING) mechanism [15–20]. Previous work has shown that inhibitory networks built to examine population activity in an *in vitro* hippocampal preparation manifest “sharp transitions” into coherent population activity caused by a small, permanent increase to the external drive to the network [21]. Additional studies have explored the effect of connection probabilities and cell characteristics manifested by classifications of cell excitability on inhibitory network synchrony [22, 23], and have noted that bistability between asynchronous and synchronous firing was possible [23].

Inhibitory oscillations are implicated in both physiological and pathological brain states. The sudden onset of inhibitory synchrony caused by an increase in drive from CA3 is suggested to underlie the generation of ripples associated with sharp waves in the hippocampus [24, 25]. Such data is further evidence that a transition to oscillatory dynamics can be brought about by increased external drive, as shown computationally [21, 23]. Inducing hyper-excitability in inhibitory cells might represent an analogue to this increased drive, potentially explaining why hyper-excitable systems transition to synchronous states of a seizure or an inter-ictal spike (IIS). Consistent with this suggestion is that interneuronal firing increases before pyramidal cell firing prior to IIS generation and seizures in animal epilepsy models [1, 26–30]. Analogously in humans, putative interneurons increase their firing before seizure onset [13].

The existing computational insights into inhibitory network synchrony, combined with the experimental literature implicating this dynamic in seizure initiation, motivate this computational study. To explore the role of interneuronal synchrony in seizure initiation, randomly connected, purely inhibitory network models are developed. These networks utilized cell models mimicking properties exhibited by neurons treated with 4-Aminopyridine (4-AP), a commonly used experimental model to generate seizures [31, 32], or properties of a healthy, control interneuron. Utilizing these tools, this investigation articulates a mechanism explaining how a sudden transition from asynchronous to synchronous firing might arise in an inhibitory network that also offers an explanation for the predisposition of hyper-excitable networks towards this transition.

This mechanism was uncovered by comparing the tendency of control and 4-AP inhibitory networks to synchronize, both from random initial conditions and following a perturbation biasing the network towards synchrony. This revealed that 4-AP networks are much more likely to transition from asynchronous to synchronous dynamics following a perturbation, due to a significantly larger regime of network parameters supporting bistability. The existence of a “bistable transition” driving an inhibitory network into synchrony expands upon existing literature probing such mechanisms, especially in the context of epilepsy. As control models do not exhibit the same predisposition for “bistable transitions” when compared to 4-AP networks, this finding may provide important insight into mechanisms underlying seizure generation. In turn, these findings provide paramount *in silico* support for the integral role of inhibitory interneurons in seizure initiation.

## Materials and methods

A link between the activity of inhibitory interneurons and seizure generation exists, although how it manifests remains unclear. Using optogenetic mice expressing channelrhodopsin-2 in inhibitory interneurons under proconvulsant conditions of 4-AP [33], it has been shown that the activation of inhibitory interneurons in layer 2-3 (L2/3) of mice somatosensory cortex can trigger ictal events [12]. The strategy here involved building generic inhibitory networks that roughly approximate cortical inhibitory networks, utilizing neuron models of both a healthy, control interneuron and an interneuron made hyper-excitable by treatment with 4-AP. Such an undertaking was informed by a combination of existing computational models of inhibitory interneurons, literature describing the general effects of 4-AP, and unpublished in-house experiments yielding data from the same interneuron in both control and 4-AP settings.

### Neuron Models

Neurons were modeled via a two dimensional system of ordinary differential equations first described by Izhikevich [34]. This model has two variables: *V*, which represents the membrane potential in mV; and *u*, which represents the slow “recovery” current in pA. The model utilized here is slightly altered in the fashion described by Ferguson et. al. [21], and is given by:

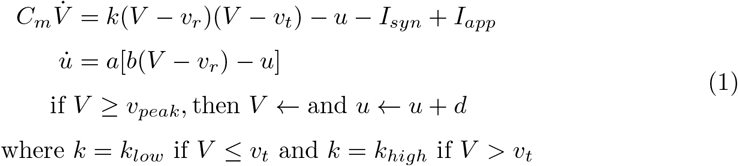

In the above equations, *C_m_* represents the membrane capacitance in pF, *v_r_* represents the resting membrane potential in mV, *v_t_* represents the instantaneous threshold potential in mV, *v_peak_* is the spike cut-off value in mV, *I_syn_* is sum of all incoming synaptic current to the neuron in pA (described in detail below), *I_app_* represents the external applied current in pA (described in detail below), *a* is the recovery time constant of the adaptation current in ms^-1^, *b* describes the sensitivity of the adaptation current to subthreshold fluctuations in nS, *c* is the voltage reset value in mV, *d* is the total current affecting the after spike behavior in pA, and *k_low_* and *k_high_* are scaling factors in nS/mV.

The use of Izhikevich model neurons was motivated by the goals of this study: namely, here we do not strictly constrain our neuron model with experimental results, but rather create a model that more “generally” matches the properties of an interneuron in both control and 4-AP cortical settings and highlights the key differences between them (particularly those caused by hyper-excitability in 4-AP interneurons). This choice allows for the detailed investigation of the mechanisms underlying the transition into synchrony in these networks performed here.

### Neuron model parameters

Models and parameter values were based primarily on previous Izhikevich inhibitory cell models [21] and the literature describing the effects of 4-AP [32]. Unpublished in-house experiments were used to supplement this literature and inform the modeling in areas in which this literature was not as detailed. These experiments highlighted specific differences in control and 4-AP settings, particularly with regards to the rheobase and capacitance.

The model presented by Ferguson et. al. [21] was used as a “starting point” for the models presented here, as the neurons of interest in that study exhibit similar major properties to the types of neurons of interest in this research. This choice informed the values of *v_r_*, *v_t_*, *c* and *v_peak_*. The unpublished experimental work yielded *C_m_* values for cortical interneurons.

The rest of the parameter values (*a, b, d, k_low_, k_high_*) were chosen through a parameter exploration to match the difference in rheobase caused by treatment of 4-AP. Unpublished in-house experiments were used for the rheobase values of control and 4-AP interneurons, as recorded in the same cell, given that such details are not available in the existing literature. An increase in spike-frequency adaptation in the 4-AP setting is also implied by the literature [32] and correspondingly influenced the determination of these parameters. As the model of Ferguson et. al. [21] was used as a reasonable model of a fast-firing inhibitory cell, the slope of the frequency-current (FI) of that neuron was used for the control case. Except for the changes caused by a shifted rheobase and the presence of adaptation, this slope was kept approximately the same for the 4-AP model. With the different rheobases, this means that the firing frequency is larger in the 4-AP model relative to control for a given input current.

The parameter values for both what will hereafter be referred to as the “control” model and what will hereafter be referred to as the “4-AP” model are included in Table 1, alongside the primary motivating factors in the choice of said parameter. PropertieS of these model neurons encapsulated in their FI curves are illustrated in Fig 2. All modeled neurons referred to as “control” or “4-AP” in this work use these parameter values (i.e. every neuron within a given network is identical with the exception of its external driving current).

**Table 1.** Parameters used in neuron models.

| Parameter | Value<br>(Control) | Value<br>(4-AP) | Rationale |
| --- | --- | --- | --- |
| $C_m$ | 73 pF | 49 pF | Unpublished in-house experiment |
| $v_r$ | -60.6 mV | -60.6 mV | Ferguson et. al. (2013) [21] |
| $v_t$ | -43.1 mV | -43.1 mV | Ferguson et. al. (2013) [21] |
| $v_{peak}$ | 2.5 mV | 2.5 mV | Ferguson et. al. (2013) [21] |
| $a$ | 0.01 ms<sup>-1</sup> | 0.01 ms<sup>-1</sup> | Parameter influences rheobase and adaptation exhibited by model* |
| $b$ | -0.2 nS | -0.4 nS | Parameter influences rheobase and adaptation exhibited by model* |
| $c$ | -67 mV | -67 mV | Ferguson et. al. (2013) [21] |
| $d$ | 1.25 pA | 0.75 pA | Parameter influences rheobase and adaptation exhibited by model* |
| $k_{low}$ | 0.4 nS/mV | 0.6 nS/mV | Parameter influences rheobase and adaptation exhibited by model* |
| $k_{high}$ | 2 nS/mV | 2 nS/mV | Parameter influences rheobase and adaptation exhibited by model* |

\* Differences in rheobase and adaptation in control and 4-AP neurons are features shown by Williams and Hablitz (2015) [32] as well as observed in our unpublished in-house experiment

**Fig 2.**
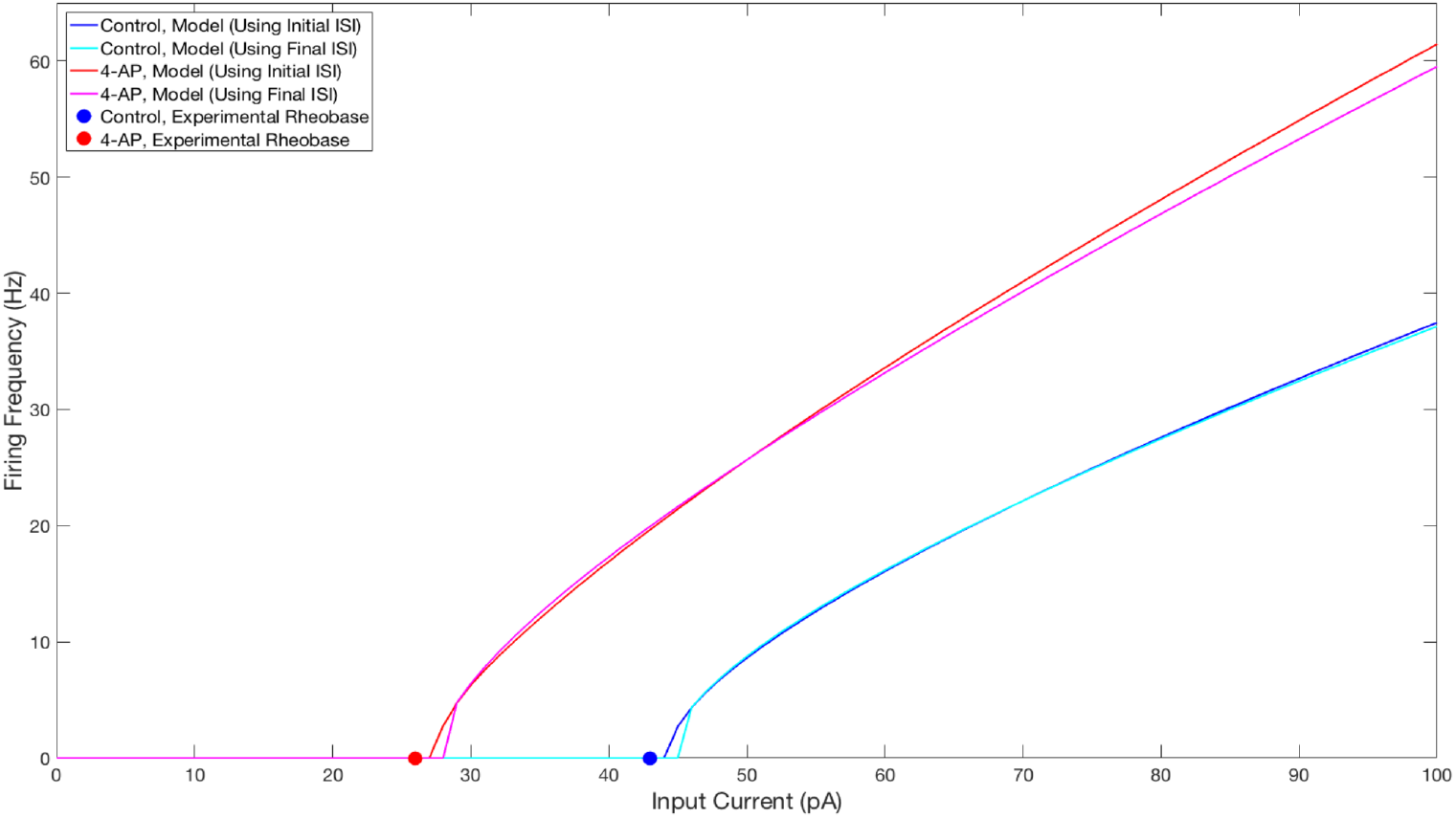
FI curves illustrating properties of neuron models used in this study. FI Curves for control (blue and cyan) and 4-AP (red and magenta) modeled neurons. Curves are shown for frequencies calculated using the initial (blue and red) and final (cyan and magenta) inter-spike intervals to illustrate the tendency for spike-frequency adaptation (SFA). These comparisons show that the neuron models utilized in this study match the decreased rheobase and increased excitability and SFA of 4-AP treated neurons in comparison to control neurons (with the rheobases determined from unpublished in-house experiment for control and 4-AP neurons highlighted on the figure by the colored dots).

### Network Structure

Similar to inhibitory network models developed by Ferguson et. al. [21], the neurons in the networks modeled here were randomly connected by synapses utilizing a first-order kinetic model. Each synapse is modeled by

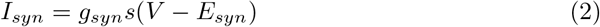

where *g_syn_* is the maximum inhibitory synaptic conductance in nS, *s* is the gating variable, *V* is the membrane potential of the post-synaptic cell in mV, and *E_syn_* is the inhibitory reversal potential in mV. As this value of *E_syn_* is set at an inhibitory value of −75 mV for every possible synapse, this study includes *only* inhibitory synaptic connections. Furthermore, *g_syn_* is uniform for each network studied, meaning each connection in a given network has the same strength.

The gating variable models the proportion of open synaptic channels, with its dynamics given by

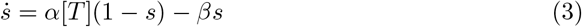

where *α* represents the inverse of the rise time constant and *β* represents the inverse oi the decay time constant [35]. [*T*] models the concentration of neurotransmitter released following a pre-synaptic action potential. [*T*] is represented as a unitary pulse lasting 1 ms, from the time of the pre-synaptic spike (*t*_0_) to the end of the pulse (*t*_1_). With this, the dynamics of *s* can be simplified to the following two equations,

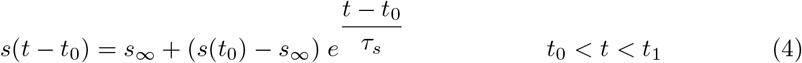

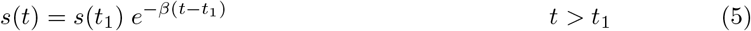

where 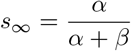 and 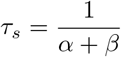.

### Network model parameters

A fast rise time rate constant of *α* = 3.7037 ms^-1^ is used here as in [21]. Values for the inhibitory reversal potential (−75 mV) and the synaptic decay rate constant (*β* = 0.3333 ms^-1^) were taken from Traub et. al. [36], and the range of inhibitory synaptic conductances explored (0 to 10 nS) encompasses cortical estimates [36].

Network size and connectivity were based on estimates regarding the density of inhibitory cells present in the cortex and their intra-connectivity [37]. Choices regarding network size were motivated by the size of L2/3 slices obtained in-house (approximately 0.03 mm^3^) combined with observations regarding the number of large basket cells (the most abundant type of inhibitory cell in L2/3) per unit volume presented in Markram et. al. [37]. Given that a single large basket cell synapses onto approximately 23 other large basket cells in this brain region [37], and assuming random connectivity, the probability of connection between such cells would be at least 0.04. Considering networks with connectivity densities lower than 0.04 would be unlikely to exhibit coherent dynamics in a randomly connected inhibitory network [23, 38], this density is used as a lower bound for this investigation. Based on such estimations, this study utilized networks of 500 neurons with the neurons randomly connected with connection probabilities of 0.04, 0.08, 0.12 and 0.16. Connection probabilities much larger than this were not needed as they would be clearly unrealistic relative to the biological estimations.

As done in Ferguson et. al. [21], cell heterogeneity in the networks was implemented by varying the amplitude of the tonic external input current, *I_app_*, to each neuron. The input currents were selected from a normal distribution with a mean value of *I_μ_*, with the degree of heterogeneity in the input currents determined by the standard deviation, *σ*. *σ* = 3, 6 and 12 pA were studied.

### Simulations

The code underlying these simulations was written in the C programming language and run on a Linux-based high-performance computing cluster utilizing Compute Canada resources provided via the University of Toronto [39]. All simulations were run for 2000 ms, with the initial conditions randomized such that *V* ∈ (−70, 0) while *u* = 0. Model equations were integrated using the Euler Method with a time step *dt* = 0.01 ms. Spikes did not trigger synaptic current until 100 milliseconds into the simulation (via a simple manipulation in the code) to allow initial transients to decay.

In order to uncover other potential dynamical states of the network, a brief, large amplitude current pulse was delivered uniformly to each cell in the network to perturb the system and potentially bias it towards the synchronous dynamical state. This 2 ms pulse had an amplitude of 1000 pA and was delivered at 1000 ms. This is analogous to imposing homogeneous initial conditions causing instantaneous spiking of all neurons in the network, in contrast to the randomized initial conditions that begin the simulations. To identify networks that exhibited bistability between asynchronous and clustered behavior, network dynamics established from random initial conditions (figure panels denoted *Random Initial Conditions*) and those established after the perturbation (figure panels denoted *Following Perturbation*) were compared.

Heatmaps of the Synchrony Measure and differences in the Synchrony Measure before and after the perturbation shown in all figures display the average of these scores over five independent simulations. The Random Initial Conditions scores were calculated based on the network activity from 500 to 1000 milliseconds, and the Following Perturbation scores were calculated based on the network activity from 1500 to 2000 milliseconds. In the heatmap plots the mean applied current value *I_μ_* was varied along the y-axis, while the inhibitory synaptic weight *g_syn_* was varied along the x-axis. Simulations (not shown here) were run to ensure that the behaviors indicated by the Synchrony Measure taken over the given intervals were indicative of stable behaviors that would persist long past the time interval measured here.

### Measures

The measure used to quantify coherent activity in the simulated networks, here termed a *Synchrony Measure*, is a slight adaptation of a commonly used measure created by Golomb and Rinzel [40, 41] that quantifies the degree of spiking coincidence in the network. This particular implementation of this measure has been utilized in previous studies [23, 42, 43]

Briefly, the measure involved convolving a Gaussian function with the time of each action potential for every cell to generate functions *V_i_*(*t*). The population averaged voltage *V*(*t*) was then defined as 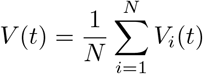, where *N* is the number of cells in the network. The overall variance of the population averaged voltage *σ* and the variance of an individual neuron’s voltage *σ_i_* were defined as

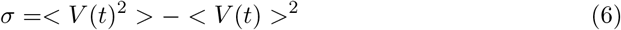

and

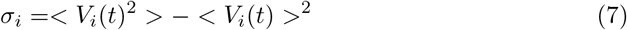

where < · > indicates time averaging over the interval for which the measure is taken. The Synchrony Measure *S* was then defined as

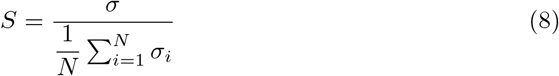

The value *S* = 0 indicates completely asynchronous firing, while *S* =1 corresponds to fully synchronous pattern of network activity. Example raster plots and the corresponding Synchrony Measure values over an illustrative range are shown in Fig 3.

**Fig 3.**
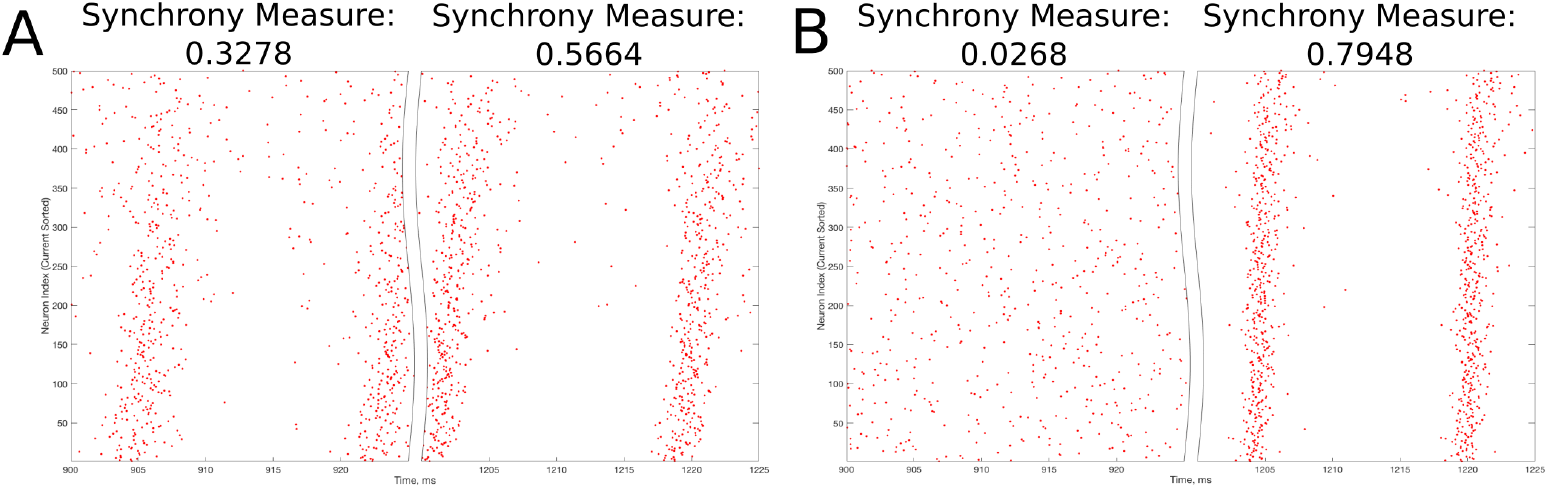
Example raster plots illustrating the Synchrony Measure and motivating the definition of the Bistability Measure. A: An example network that exhibits “messy” synchronous dynamics both before and after the perturbation is delivered at 1000 ms, resulting in a moderate value of the Synchrony Measure in each case. Dynamics before the perturbation are shown in the left panel, while dynamics following the perturbation are shown in the right. Although the Synchrony Measure following the perturbation is larger than that before the perturbation, this increase does not indicate a bistable transition from asynchronous to synchronous dynamics, but rather qualitatively “tighter” synchrony. The choice of 0.3 as the “threshold value” in the articulation of the Bistability Measure prevents cases such as this from contributing positively to the measure. B: An example network exhibiting asynchrony before the perturbation (left panel) and very clear synchrony afterwards (right panel), along with the corresponding Synchrony Measures. Very low synchrony measures (typically less than 0.25) indicate asynchrony, while higher Synchrony Measures illustrate synchrony, with higher values indicating more ordered and less “messy” synchrony.

This research was interested not merely in the degree of synchronous firing in the networks of interest (described by the Synchrony Measure), but rather was primarily focused on identifying a transition from asynchronous to synchronous dynamics driven by network bistability. A straightforward way to identify whether such a transition occurred following a perturbation (discussed in detail above) is to compare the value of the Synchrony Measure before and after said perturbation; in such a comparison, large increases in the Synchrony Measure following the perturbation are likely indicative of a transition from asynchronous to synchronous dynamics. Further, to analyze a network’s predisposition towards this transition over a range of parameter values, one need only summate these individual comparisons in an informed fashion into a single, quantitative score. This motivated the creation of a *Bistability Measure* used in this study.

This measure was calculated in three steps. First, the difference between the Synchrony Measure following the modeled perturbation and the Synchrony Measure from randomized initial conditions was taken for each network in some context (i.e. for a particular parameter range encapsulated by a given heatmap). Second, only cases when this difference was > 0.3 were included in the summation (this choice is justified in detail below). Finally, the sum of the Synchrony Measure differences exceeding this threshold value was taken to yield the final Bistability Measure.

The choice of the “threshold” value of 0.3 in the second step above merits further explanation. The Synchrony Measure is not a binary differentiation between asynchronous and synchronous dynamics, but rather a quantitative measure of the degree of synchronous firing. This means that increases in the Synchrony Measure, particularly subtle ones, do not necessarily indicate a differentiation of asynchrony from synchrony, but instead could indicate the presence of qualitatively “tighter” synchrony. An example of such a case is seen in Fig 3A. However, large increases in the Synchrony Measure are almost always indicative of entirely different dynamical states, as shown by the example in Fig 3B). After a thorough investigation of the correspondence between a qualitative assessment of synchrony (i.e. visual inspection of raster plots) and the quantitative assessment provided by the Synchrony Measure, it was determined that a difference of at least 0.3 in the Synchrony Measure before and after a perturbation best identified networks in which a transition between dynamical states occurred while excluding networks in which an increased Synchrony Measure only indicated subtle changes in the network dynamics.

Instantiating this “threshold” value into the calculation of the Bistability Measure ensures that the measure best quantifies the tendency for networks to exhibit bistable transitions, rather than naively quantifying the difference in Synchrony Measure before and after the perturbation. This occurs in two fashions during the calculation of the measure to further ensure robustness: first, networks that exhibit minor changes in the Synchrony Measure (< 0.3) are completely excluded from the summation, considering such networks are extremely unlikely to exhibit a bistable transition; and second, the summation of the change in the Synchrony Measure values, rather than a binary summation of which networks exhibit a change above the threshold value, allows networks that exhibit a larger Synchrony Measure difference (for which one can much more confidently assert a dynamical transition occurs) to be weighted more heavily in the calculation of the Bistability Measure. Finally, the fact that this value was not chosen arbitrarily is worth further emphasis: this choice was made only after a detailed investigation into the interpretation of various Synchrony Measure differences and trial calculations of the Bistability Measure with various choices of this “threshold” value (not shown here), all of which that contained flaws improved upon by the choice made for the final measure.

### Ornstein-Uhlenbeck process

The perturbation described above is motivated primarily by the desire to uncover a mechanism for the transition from asynchrony to synchrony from the perspective of dynamical systems. In order to assess whether this mechanism is biologically reasonable, an analogous perturbation that might arise in more biologically grounded models was sought.

An Ornstein-Uhlenbeck process [44] is used in the literature to model background synaptic input into a network [45, 46], and is used in this study to determine whether “perturbation-like” activity might arise naturally from this model of external synaptic input. This process, used to determine the conductance of excitatory synaptic input in this context, is described mathematically by the following equations [45] with an initial condition *g_e_*(0) = *g*_*e*(0)_:

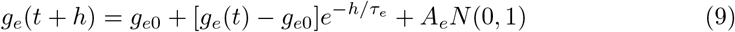

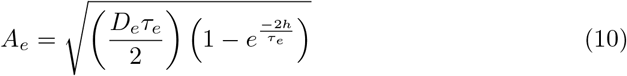

where *N*(0, 1) is a normal random number taken from a distribution with 0 mean and a standard deviation of 1.

The insights from Piwkowska et. al. [46] allowed for the choice of parameters constrained by cortical data. The parameters used in the Ornstein-Uhlenbeck process utilized in this study were *g*_*e*0_ = 3 nS, *τ_e_* = 2 ms, and *D_e_* = 2 (a unitless diffusion coefficient), and the integration time step was *h* = 0.01 ms.

### Code Accessibility

The code/software described in the paper is freely available online at https://github.com/FKSkinnerLab/CorticalInhibitoryNetwork.

## Results

In order for inhibitory interneurons to serve as the impetus for seizures, as posited by a “GABAergic initiation hypothesis”, there should exist a mechanism that explains both how a network of such interneurons can suddenly transition into synchronous dynamics (which in turn provides a large burst of synaptic inhibition to pyramidal neurons in the behaving animal) and why model seizure networks are more prone to exhibit this transition and the resultant pathological brain activity. To examine this, an experimental 4-AP hyper-excitable state, as used to model seizures by Chang et. al. [12], is used as a basis for these mathematical model explorations, with results from such networks compared to model control networks.

### Bistability between coherent and incoherent states is exhibited more robustly by 4-AP inhibitory networks

The concept of bistability arises primarily from the mathematical study of dynamical systems. In this context, a “stable” state is one which will be preserved by the system for all time in the absence of any perturbations to the conditions defining the system. In nonlinear systems it is possible for multiple stable states to exist, and for the network to naturally settle into any one of these stable states depending upon a variety of factors including the initial conditions and any perturbations that might be delivered. In a biological system, this could manifest from the history of inputs from different brain structures along various pathways to the network in question. Such a system is defined to be “bistable” or “multistable” given the existence of more than one stable solution to the mathematical equations [47].

The results presented in Fig 4 show that many of the networks within the parameter regime considered in this work exhibit bistability. In panels A and B the Synchrony Measure (described in the Materials and Methods section) was taken for the same networks in two different states: the results from randomized initial conditions are shown in the left panels, while the results following a perturbation to the system (described in the Materials and Methods section) are shown in the right panels. Note that the parameter range shown in these heatmaps is “zoomed in” relative to the larger parameter scan used in the heatmaps presented in the following section in order to better highlight the regime of bistability. Control networks are shown in panel A while 4-AP networks are shown in panel B.

**Fig 4.**
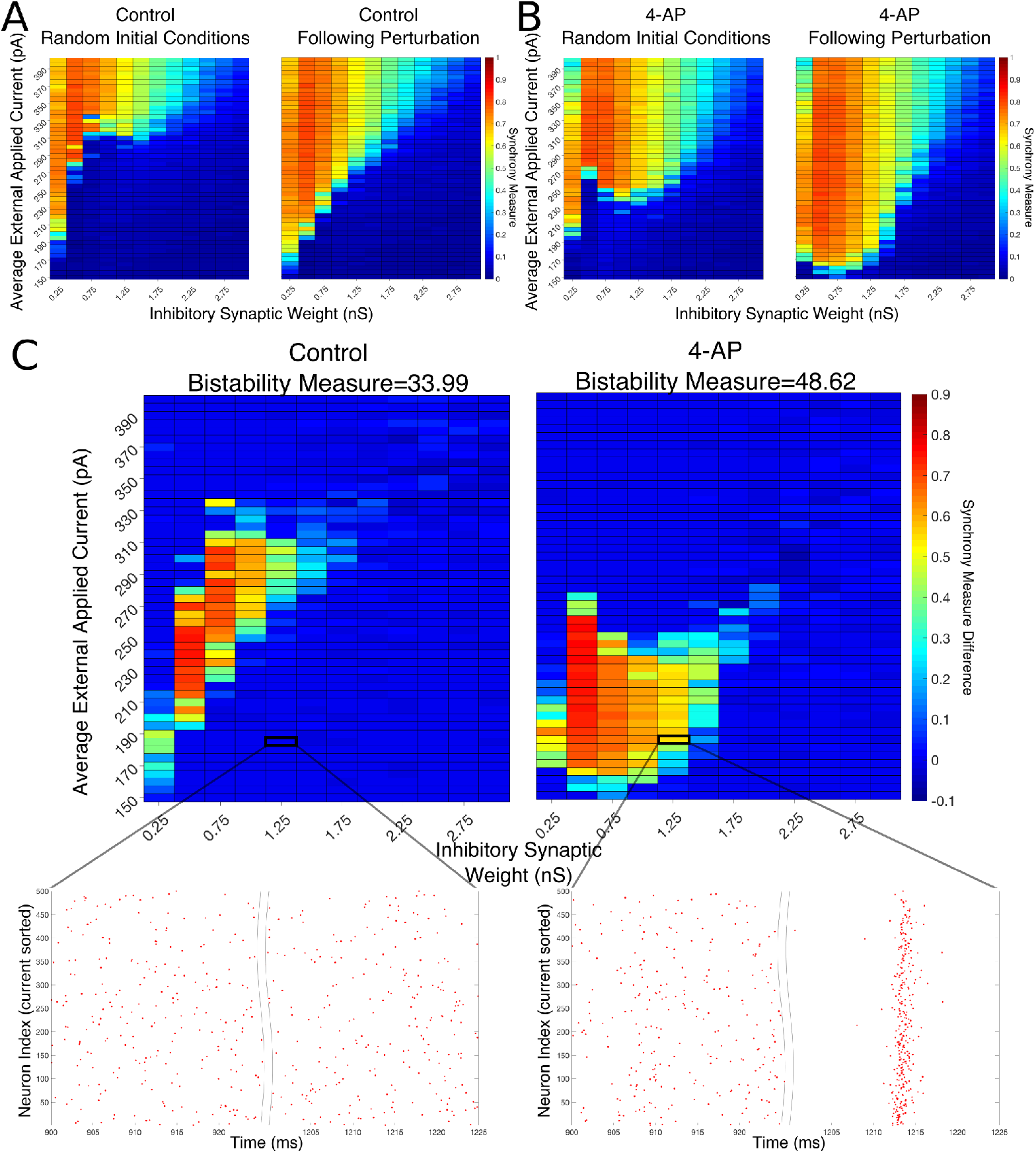
Networks containing model 4-AP neurons are more prone to bistability than networks of control neurons. A-B: Heatmaps displaying the Synchrony Measure for control networks (A) and 4-AP networks (B) with a connectivity probability of 0.12 and a standard deviation amongst the driving currents of 6 pA. In these heatmaps, the inhibitory synaptic weight is varied along the x-axis and the average external applied current is varied along the y-axis. The left panel displays the measure taken from randomized initial conditions, while the right panel displays the measure taken after a modeled perturbation. C: Heatmaps over the same parameter regime, but now showing the difference between the Synchrony Measure shown in the right and left heatmaps in panels A and B. Control results are shown on the left, and 4-AP results are shown on the right. 4-AP model networks are much more likely than control model networks to exhibit a change in dynamics following the perturbation (as indicated both by more warm colors in the heatmap and by the increased Bistability Measure score shown above the panels), indicating that the perturbation induced a transition from asynchronous to synchronous dynamics indicative of a bistability. A raster plot for both the control and 4-AP settings for a network with an inhibitory synaptic weight of 1.25 nS and an average external applied current of 185 pA (corresponding to the outlined box in the heatmap) is shown, providing an illustrative example of a case where the transition from asynchronous to synchronous dynamics following the perturbation, and thus the existence of a bistability, is observed in the 4-AP but not the control case.

There appear to be a number of networks in both the control and 4-AP settings that show a high Synchrony Measure, and thus coherent network states, following the perturbation but not from randomized initial conditions. This is indicative of a bistable system in which both the coherent and incoherent states are stable, even though the network might require a perturbation in order to leave the incoherent stable state and settle into the coherent stable state. This result is highlighted by Fig 4C in which the difference between the Synchrony Measure following the perturbation and the Synchrony Measure from randomized initial conditions is plotted to highlight the networks in which this difference occurs. Qualitatively, it appears not only that the parameter regime including these type of networks is shifted when comparing the 4-AP and control cases, but most importantly it appears that *more* of these types of networks exist in the 4-AP setting as opposed to the control case. To quantify this observation, a Bistability Measure (as outlined in the Materials and Methods section) was used, revealing that, indeed, the parameter regime defining bistable networks is larger in the 4-AP case. Raster plots highlighting an example network that is bistable in a 4-AP network, but not in the control case, for the same parameter values are shown below these heatmaps.

It makes sense, in the context of the study of seizure initiation, that both control and 4-AP networks would exhibit some bistability. Indeed, it is well established that all brains are capable of generating a seizure, even though seizures are much more likely in individuals with epilepsy (see the literature on seizures arising in non-epileptic patients following traumatic brain injury [48, 49]). However, it is interesting in the context of a “GABAergic initiation hypothesis” that 4-AP networks were more likely to exhibit bistability than control networks. This result supports the hypothesis that 4-AP treated, hyper-excitable interneurons are more likely to exhibit the necessary dynamics underlying seizure onset, which in this context is the transition from asynchronous to synchronous firing via a “bistable transition”. It is also interesting to note that the bistable regime is both wider (i.e. encompassing a larger range of synaptic strengths) and includes lower driving currents for 4-AP networks, although the latter is perhaps expected due to the lower rheobase of 4-AP neurons.

The robustness of this result was confirmed when networks were subjected to different degrees of heterogeneity in the external driving currents and different connection probabilities. This is shown by the Synchrony Measure difference heatmaps and Bistability Measures shown in Fig 5. Indeed, in all four cases presented (varying connection probability in panels A-B and varying standard deviation of the external applied currents in panels C-D, the 4-AP networks were more likely to exhibit bistability, as seen via a joint analysis of the Bistability Measures and the bistable parameter regime in the heatmaps.

The analysis of these *in silico* networks through the lens of the mathematical concept of bistability reveals crucial properties of 4-AP networks that could not otherwise be identified. However, the question remains whether a transition of this type is biologically feasible, especially considering the perturbation used to reveal the existence of the bistability was motivated from dynamical systems insights rather than the underlying biology. We address this using an Ornstein-Uhlenbeck process (as described in the Materials and Methods section [45, 46]) to generate a reasonable approximation of background excitatory synaptic conductance in the cortex. Such synaptic activity can be thought of as a more biologically-grounded analogue for the *I_app_* tonic driving current used in the computational models here. The conductance generated by the Ornstein-Uhlenbeck process is transformed into a driving current simply by multiplying by (*V* − *E_syn_*), where here *E_syn_* takes on an excitatory value of 0 mV and V is approximated as the resting potential of the neuron (here −60.6 mV).

**Fig 5.**
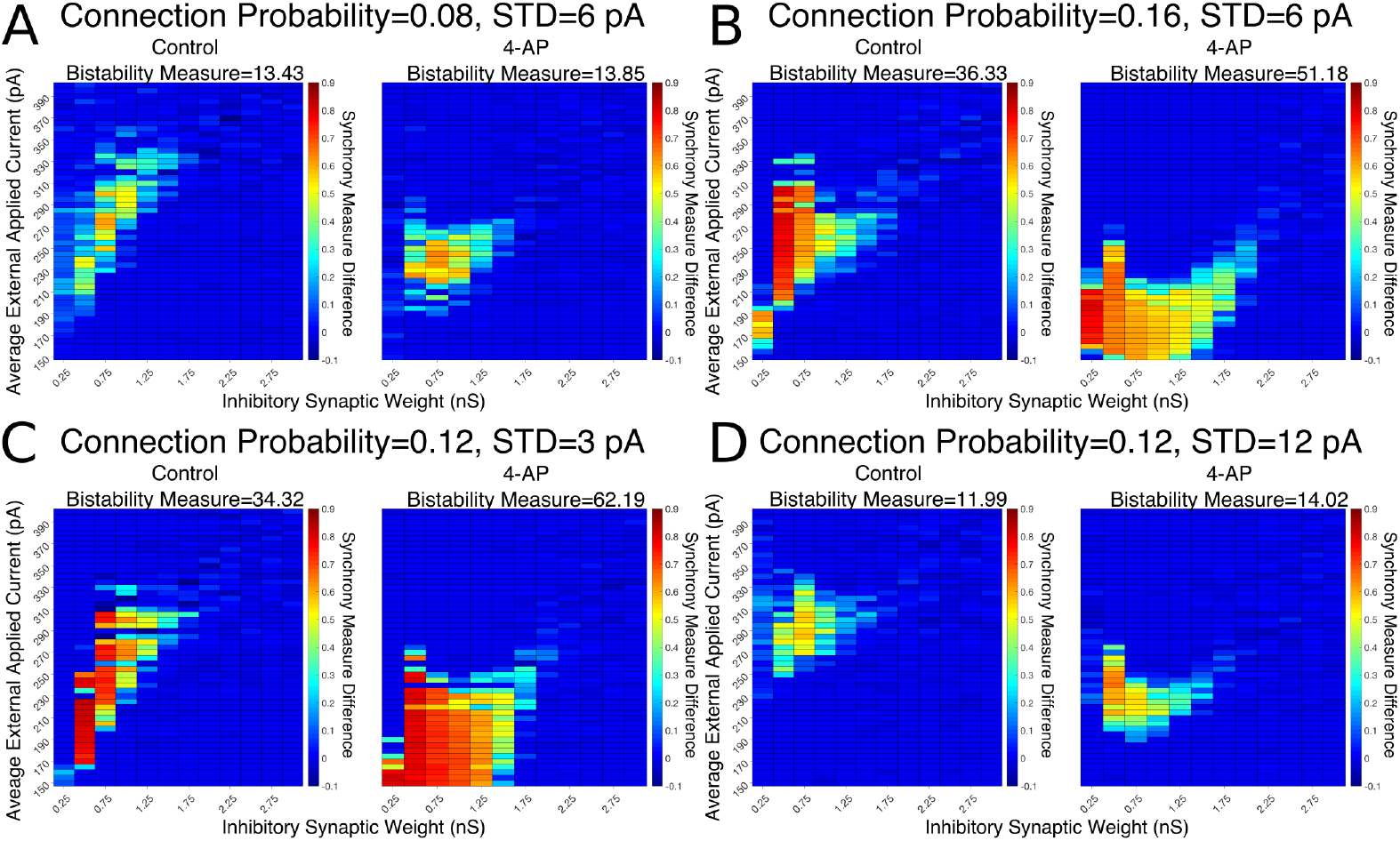
Networks containing model 4-AP neurons exhibit bistability more robustly than control networks for a variety of network parameters. A-D: Heatmaps displaying the difference between the Synchrony Measure following a modeled perturbation and from randomized initial conditions for control networks (left) and 4-AP networks (right). In these heatmaps, the inhibitory synaptic weight is varied along the x-axis and the average external applied current is varied along the y-axis. The Bistability Measure for each condition is shown above the corresponding panel. Results for a connection probability of 0.08 and standard deviation of 6 pA are shown in panel A, results for a connection probability of 0.16 and standard deviation of 6 pA are shown in panel B, results for a connection probability of 0.12 and standard deviation of 3 pA are shown in panel C, and results for a connection probability of 0.12 and standard deviation of 12 pA are shown in panel D. In all cases, 4-AP model networks exhibit a larger parameter regime showing behaviors indicative of a bistability than analogous control networks, shown both by more warm colors in the heatmap and the increased Bistability Measure.

An example of such a current, generated for 1000 ms, is seen in Fig 6A. Zooming in on the red portion of the current (225 ms to 275 ms), a 5 ms portion of the current trace that retains a significantly higher than average value is highlighted in green. This current is simplified for computational implementation by a square current pulse with an amplitude of 320 pA and a 5 ms duration, approximated on the figure with a dotted black line.

**Fig 6.**
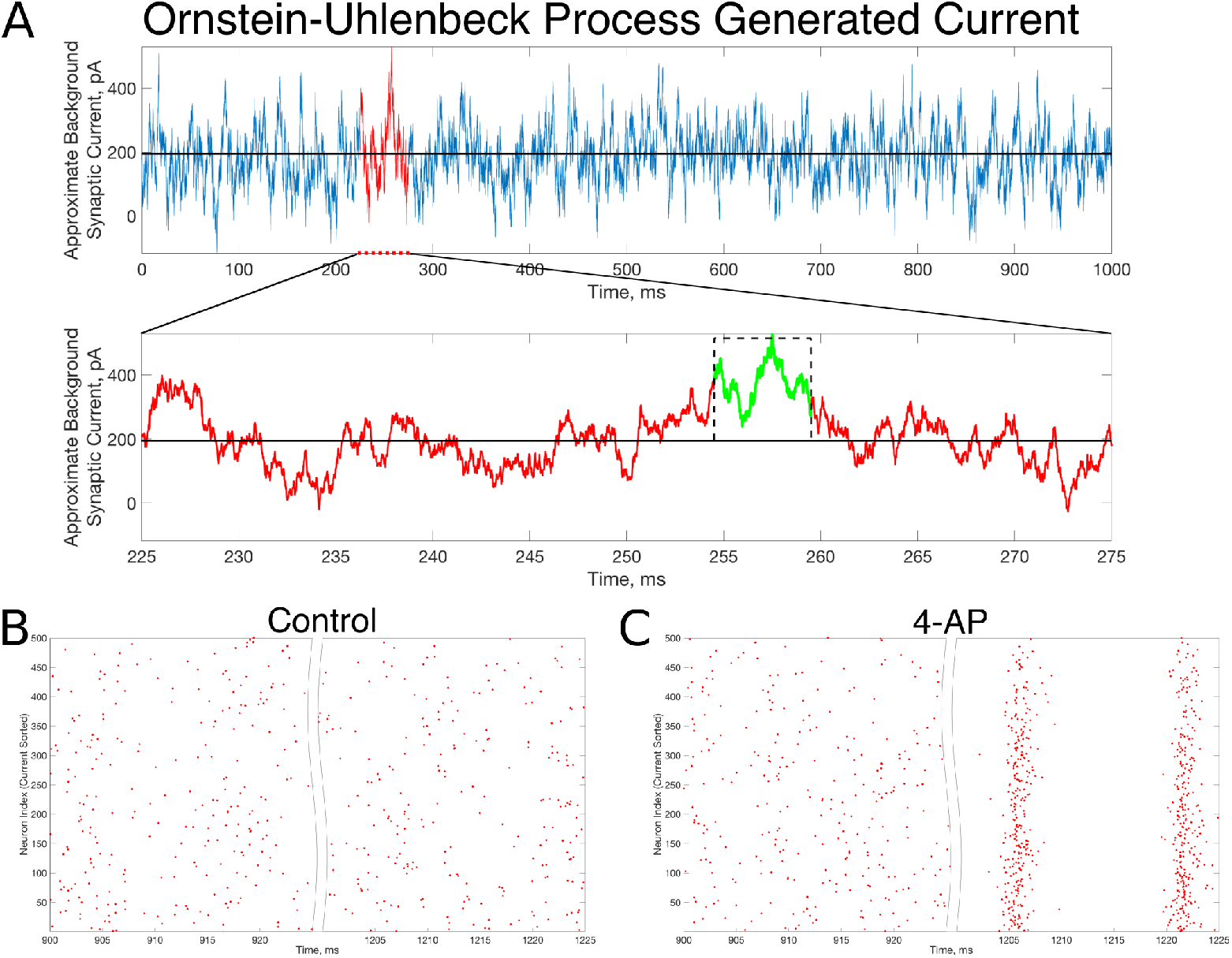
In *vivo*-like excitatory background synaptic currents can also elicit “bistable transitions” in the model inhibitory networks. A: An example of excitatory background synaptic current (1000 ms in the top panel) generated using an Ornstein-Uhlenbeck Process with parameters informed by cortical experimental literature. The bottom panel zooms in on a region of interest (plotted in red) revealing a brief period (plotted in green) in which the current is significantly larger than its average value, activity which has perturbation-like qualities. This activity is approximated by a current pulse of similar width and amplitude, plotted on the figure in a dashed black line. B-C: Raster plots for a control (B) and 4-AP (C) network that is identical to the examples displayed in Fig 4, where the large, brief current pulse used as the perturbation throughout this study is replaced by a current pulse informed by the Ornstein-Uhlenbeck Process shown in panel A that represents *in vivo*-like activity. Despite this change, which amounts to a wider pulse with significantly lower amplitude, the control and 4-AP networks still exhibit antithetical responses to this perturbation; namely, control networks return to asynchronous firing following the perturbation while 4-AP networks transition into synchronous dynamics.

Critically, the “bistable transition” typified by the raster plots in Fig 4 is preserved when the perturbation is replaced by the current pulse motivated by the results from the Ornstein-Uhlenbeck process, as shown in Fig 6B-C. This result suggests that a “bistable transition” is viable in a more biologically-grounded setting, as it can be triggered by a perturbation that could reasonably occur due to fluctuations in the background excitatory synaptic activity in the cortex. Taken together with the detailed analysis presented above of the bistability present in these networks from the perspective of dynamical systems, it is apparent that a transition from asynchrony to synchrony in inhibitory networks caused by a “bistable transition” is both a computationally and biologically plausible mechanism explaining the initial step of a “GABAergic initiation hypothesis” of seizure.

### Sharp transitions between coherent and incoherent states caused by increased external input are not likely to underlie the initial step of seizure initiation

In inhibitory network models of CA1 hippocampus that were constrained in size, connection probability, cellular and synaptic properties, previous work by [21] demonstrated “sharp transitions” between asynchronous and synchronous firing caused by a small, permanent increase in the external drive to the network. This additional mechanism has both experimental and computational support (see the discussion in the Introduction) to explain a transition into synchrony in purely inhibitory networks. We thus investigated it as a potential mechanism for the initial inhibitory synchrony necessary for a “GABAergic initiation hypothesis” of seizure. Indeed, potentially eliminating it as a candidate mechanism in this context would provide additional support for the viability of the “bistable transition” mechanism described in the previous section.

We investigated the tendency for inhibitory networks of both control and 4-AP neurons to synchronize from randomized initial conditions with varying connection probabilities and heterogeneities. Fig 7 shows results illustrating network coherence for a parameter scan over a range of inhibitory synaptic strengths that encompass physiological estimates [37] and average external applied currents with varied connection probabilities. Panels A-C show Synchrony Measure scores for networks over the entire parameter regime studied here, with results for control networks shown on the left and 4-AP networks shown on the right.

**Fig 7.**
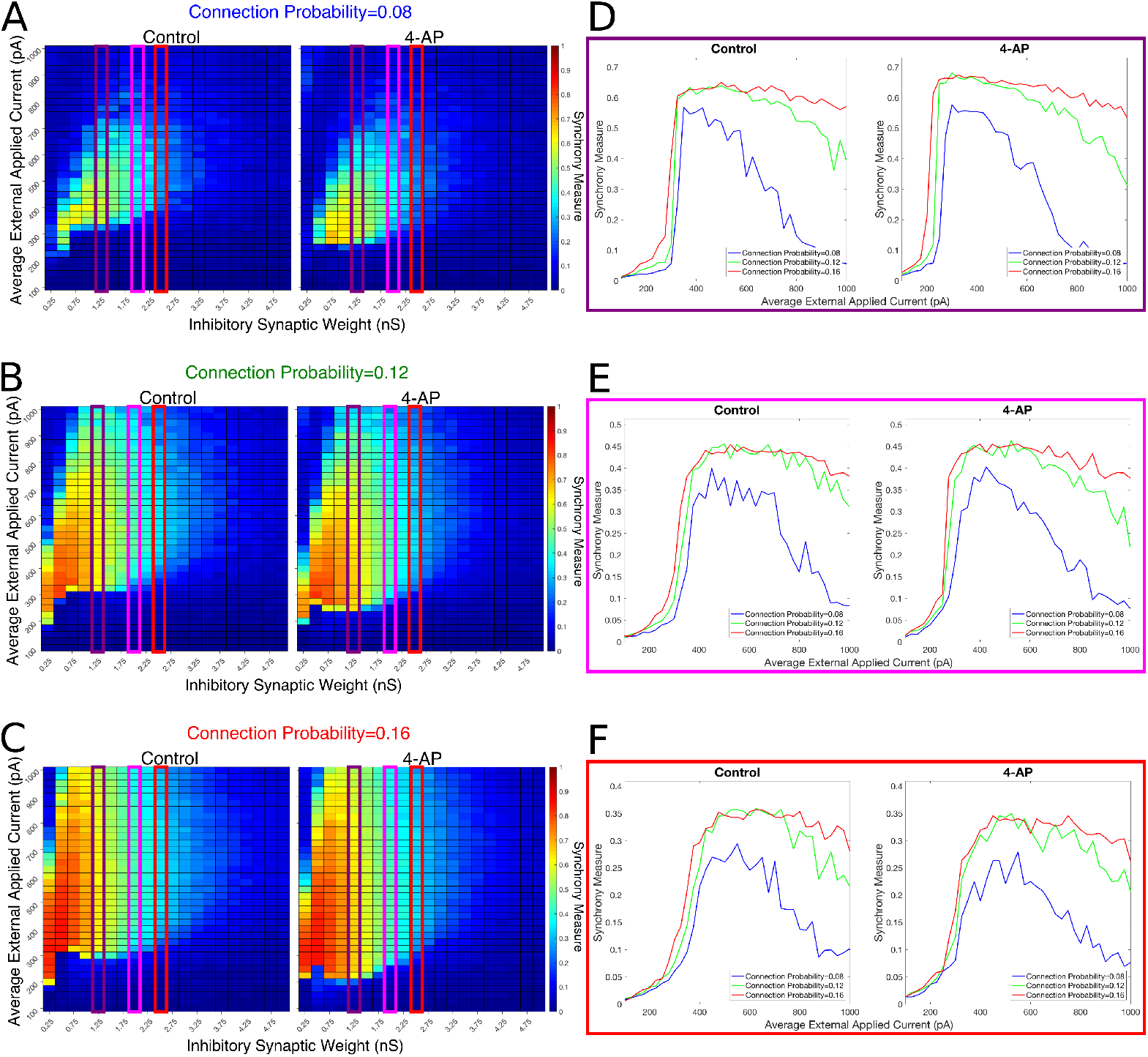
Cortically motivated inhibitory networks exhibit a “sharp transition” between asynchronous and synchronous dynamics driven by an increase in the external drive for various connection probabilities. A-C: Heatmaps displaying the Synchrony Measure for control networks (left) and 4-AP networks (right) with a standard deviation amongst the driving currents of 6 pA and varying connectivity densities. In these heatmaps, the inhibitory synaptic weight is varied along the x-axis and the average external applied current is varied along the y-axis, and the measure is taken from random initial conditions. D-F: Two dimensional “slices” of the heatmaps in panels A-C taken to better illustrate the sharpness of the transition between asynchronous and synchronous dynamics as well as more easily compare this sharpness both across varying connection probabilities and between control and 4-AP conditions. Panel D shows results for an inhibitory synaptic weight of 1.25 nS, panel E for an inhibitory synaptic weight of 2.0 nS, and panel F for an inhibitory synaptic weight of 2.5 nS. There is no significant difference in the tendency for 4-AP versus control networks to exhibit the “sharp transition” from asynchrony to synchrony despite differences in the parameter regime supporting synchrony. Furthermore, the differences in the “sharpness” of the transition in the two cases are not robust.

While connection probability estimates indicated a value of at least 0.04 was biologically reasonable (see Methods), simulated networks produced no coherent states with this connection probability. This is perhaps not too surprising given that the cellular models utilized here were only loosely motivated by experiments (see Methods) so that additional estimates of network connectivity are not expected to be precise. However, it is expected that any differences in control and 4-AP models are meaningful since these differences were captured in a comparable fashion (see Methods and Fig 2).

Fig 7 also includes three two-dimensional plots highlighting the evolution of the Synchrony Measure as a function of the average external applied current for a set value of the inhibitory synaptic weight in panels D-F. Results for each connection probability are shown jointly to facilitate comparison, with results for control networks shown in the left panels and results for 4-AP networks shown in the right panels. Additionally, the “sharpness” of the transition from asynchrony to synchrony was quantified by taking the slope of the line segment best representing this transition, which is chosen to be that between the first point that achieves a Synchrony Measure greater than half the maximum Synchrony Measure observed by networks in that panel and the point one current step earlier. The slopes for all of the examples presented in Figs 7 and 8 are shown jointly in Table 2.

**Fig 8.**
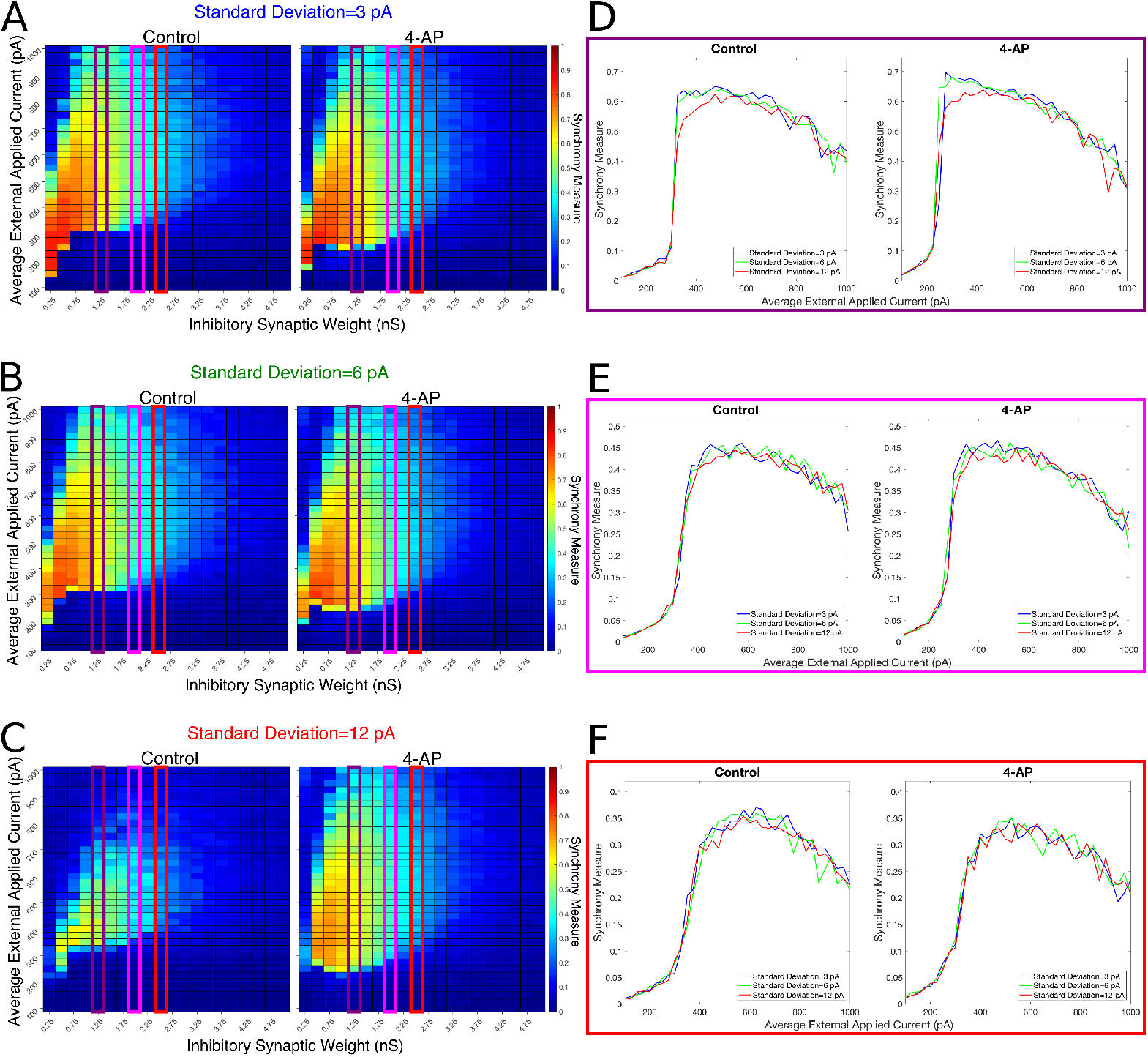
Varying the heterogeneity in external driving current in modeled purely inhibitory networks largely preserves the general dynamical differences and similarities seen between the 4-AP and control cases from randomized initial conditions. A-C: Heatmaps displaying the Synchrony Measure for control networks (left) and 4-AP networks (right) with a connection probability of 0.12 and varying standard deviations amongst the driving currents. In these heatmaps, the inhibitory synaptic weight is varied along the x-axis and the average external applied current is varied along the y-axis, and the measure is taken from random initial conditions. D-F: Two dimensional “slices” of the heatmaps in panels A-C taken to better illustrate the sharpness of the transition between asynchronous and synchronous dynamics as well as more easily compare this sharpness both across varying connection probabilities and between control and 4-AP conditions. Panel D shows results for an inhibitory synaptic weight of 1.25 nS, panel E for an inhibitory synaptic weight of 2.0 nS, and panel F for an inhibitory synaptic weight of 2.5 nS. Once again, there is no significant difference in the tendency for 4-AP versus control networks to exhibit the transition from asynchrony to synchrony, nor any significant differences in the “sharpness” of this transition.

**Table 2.** Slopes quantifying “sharpness” of the transition from asynchrony to synchrony seen in Figs 7 and 8.

| Connection Probability | Standard Deviation | Inhibitory Synaptic Weight | Control Slope | 4-AP Slope |
| --- | --- | --- | --- | --- |
| 0.08 | 6 pA | 1.25 nS | 0.0125 | **0.0135** |
|  |  | 2.00 nS | **0.0059** | 0.0047 |
|  |  | 2.50 nS | **0.0030** | 0.0012 |
| 0.12 | 3 pA | 1.25 nS | **0.0204** | 0.0172 |
|  |  | 2.00 nS | 0.0074 | **0.0096** |
|  |  | 2.50 nS | **0.0041** | 0.0033 |
| 0.12 | 6 pA | 1.25 nS | 0.0191 | **0.0211** |
|  |  | 2.00 nS | 0.0038 | **0.0057** |
|  |  | 2.50 nS | 0.0027 | **0.0042** |
| 0.12 | 12 pA | 1.25 nS | 0.0135 | **0.0138** |
|  |  | 2.00 nS | 0.0043 | **0.0074** |
|  |  | 2.50 nS | 0.0034 | **0.0039** |
| 0.16 | 6 pA | 1.25 nS | 0.0106 | **0.0162** |
|  |  | 2.00 nS | 0.0059 | **0.0108** |
|  |  | 2.50 nS | 0.0027 | **0.0037** |

Note: for each parameter set, the larger slope is bolded for ease of reference.

The results presented in Fig 7 show that a sharp transition between asynchrony and synchrony caused by a small, permanent increase in external driving current does occur in these cortically-motivated networks over a range of reasonable connection probabilities, both for control and 4-AP neurons. As one would intuitively expect, the parameter regime in which network coherence occurs grows larger as the connection probability becomes larger (Fig 7A-C. The two-dimensional plots (Fig 7D-F do not show a clear pattern between the connection probability and the sharpness of the transition, but this is reasonable considering that large connection probabilities were not included in our explorations (see Methods). The differences between control and 4-AP networks were also observed when the heterogeneity was varied as shown in Fig 8. Heatmaps analyzing the Synchrony Measure over the entire parameter regime are shown in panels A-C, with similar comparison between control and 4-AP networks as in Fig 7, while analogous two-dimensional Figs to those in Fig 7 are shown in panels D-F, but with varying standard deviations as opposed to connection probabilities in each panel. The results presented in Fig 8 show the expected effects of increased heterogeneity: namely, as the heterogeneity increases, the size of the parameter regime exhibiting coherent states decreases. This is shown most clearly by comparing the results with a standard deviation of 12 pA to both the results with a standard deviation of 3 pA and 6 pA, which show similar regimes of synchronous dynamics (although the synchrony is more pronounced over this regime when the heterogeneity is smallest at 3 pA).

While these results show the existence of transitions caused by increased external drive that are “sharp”, Figs 7 and 8 do not reveal any difference in the tendency for control or 4-AP networks to exhibit this sharp transition. While 4-AP networks exhibit a high synchrony measure over a wider parameter regime, particularly at lower values of the average external applied current (explained by a combination of the hyper-excitability of 4-AP neurons, insights from the ING mechanism, and the analysis of sparsely connected inhibitory networks presented by Rich et. al. [23]), this does not indicate an increased tendency to exhibit the *transition* from asynchrony to synchrony. Indeed, such a transition exists almost uniformly across the inhibitory synaptic weights studied here (which can be seen both by visually inspecting the increase in Synchrony Measure going up a column in the heatmaps or looking at the two-dimensional traces), with only the applied current value at which the transition occurs changing.

Furthermore, any comparisons of the relative “sharpness” of the transitions in control and 4-AP networks are qualitative at best. In a majority of the comparisons illustrated in the two-dimensional plots (Fig 7D-F), 4-AP networks displayed a higher slope measure shown in Table 2 than their control counterparts. However, this feature is not entirely robust (see, for example, the comparison of networks with a connection probability of 0.08 in Fig 7E and Table 2), and there is no guarantee that the minor increases in the slope measure are indicative of a difference in the actual dynamics underlying the transition. Further detailed analysis of this feature of the networks would be required to draw any conclusions.

Finally, it is worth noting that a change of this sort is unlikely to arise via an Ornstein-Uhlenbeck process modeling background excitatory synaptic activity, in contrast to what was shown in Fig 6 (i.e. that perturbation-like activity could arise from this process). Indeed, the example current presented in Fig 6 does not show any large amplitude increases in the synaptic current lasting longer than tens of milliseconds. This indicates that a small, permanent increase to the external drive to the network likely requires a more consequential biological change in the system, especially when compared to a perturbation which can arise more naturally via fluctuations in the background synaptic activity.

Taken together, these results confirm that a transition from asynchrony to synchrony as a result of minor, permanent increases to the external driving current can occur in these cortically-motivated networks, similar to the results presented by Ferguson et. al. [21] in the hippocampus. However, there is no robust difference in the tendency for 4-AP versus control networks to exhibit this transition. This strongly suggests that this mechanism is unlikely responsible for the initial step in a “GABAergic initiation hypothesis”. Instead, a mechanism driven by a “bistable transition” is more plausible in the context of explaining GABAergic seizure initiation, as it is much more likely to occur in 4-AP networks rather than control networks. While transitions into synchrony caused by minor, permanent increases to the external drive to an inhibitory network certainly could occur in the brain given the existing literature, this conclusion implies that mechanism is more likely to drive non-pathological oscillations rather than the pathological inhibitory synchrony potentially initiating seizure.

## Discussion

Computational models of epilepsy encompass various levels of detail and address different seizure aspects [50]. To help individuals with epilepsy, mechanisms underlying the initiation, propagation and termination of seizures must be discovered and fully understood (see Fig 1). This paper aims to provide support for a novel hypothesis for seizure initiation, a “GABAergic initiation hypothesis”, by proposing a viable cellular-based mechanism by which the necessary initial step of this hypothesis, the sudden transition of ictogenic inhibitory networks into synchrony, might come about. Such a mechanism has not yet been presented in the literature of which the authors are aware. This hypothesis proposes that synchronous activation of inhibitory interneurons (the “initial step” investigated here) gives rise to a strong inhibitory signal that activates the excitatory, pyramidal cell population via PIR [12] (see Fig 1). As experimental support for this theory accumulates [1, 8–13], computational insights such as those presented here can provide further support for its viability by proposing reasonable mechanisms underlying the activity seen in experiments. Such mechanistic insights may also facilitate future clinical applications of this research.

In this study, inhibitory networks informed by a cortical environment in control and hyper-excitable settings were constructed, with the modeled hyper-excitability specifically mimicking the treatment of interneurons with 4-AP. Experimentally, *in vivo* and *in vitro* treatment with 4-AP induces seizures that are preceded by interneuronal synchrony and predominantly GABAergic IIS [29, 51], and thus 4-AP is a commonly used epilepsy model [31, 32]. This experimental evidence justifies not only the use of 4-AP as a seizure-model in this study, but also the focus on the effects of 4-AP on interneuronal networks. However, GABAergic activity does not only play a role in seizure initiation under 4-AP conditions; indeed, synchronous interneuronal activation has also been shown to underlie IIS in the *in vivo* pilocarpine model of epilepsy [13], precede seizures in both the low-Mg, high K+ model [26, 28] and electrical stimulation models of seizure initiation [52], and more generally precede seizures in rodents [9, 29]. Thus, the insights gained from this study can be applied to the general study of epileptiform activity.

Two potential mechanisms by which inhibitory networks could suddenly transition into synchronous firing were examined in this paper. In the context of the study of seizure initiation, the mere existence of such a transition is not of primary concern; rather, such a transition should occur appreciably more often in hyper-excitable (i.e. 4-AP treated) networks when compared to control networks. One potential mechanism, previously articulated in the computational literature by Ferguson et. al. [21] in a hippocampal setting and confirmed by other studies for more general networks [23], proposes that small, permanent increases in excitatory drive could cause a sharp transition between incoherent and coherent states in a purely inhibitory network. While transitions of this type were present in the networks studied here, there was no difference in the tendency for this transition to occur when 4-AP and control networks were compared. In contrast, transitions caused by a brief perturbation to external drive to the system, termed “bistable transitions” given the correspondence of this transition with the concept of bistability from dynamical systems theory, were notably more likely to occur in 4-AP than control networks. This crucial difference implies that “bistable transitions” are a more viable candidate mechanism that explains a sudden transition of an inhibitory network into synchrony in pathological networks.

The general concept of bistability has been discussed previously in epilepsy literature, given that epilepsy as a disease represents the sudden transition between two seemingly stable brain states: the “healthy” non-seizure state characterized by largely uncorrelated neural activity and the “pathological” seizure state characterized by synchronous neural firing [53]. However, it is worth emphasizing that the *setting* in which bistability is analyzed, and thus the context of the corresponding perturbation, is unique in this study, driven primarily by the focus on a “GABAergic initiation hypothesis”. Indeed, existing studies investigate a bistability between seizure and non-seizure states in settings such as intact hippocampal slices [54], a computational network of both excitatory and inhibitory cells with special emphasis on the role of extracellular potassium concentrations [55], or more general mathematical settings [53]. In contrast, in this study bistability is analyzed solely in an inhibitory network, and the bistability does not in itself represent the transition into seizure, but rather a dynamical change that might precipitate seizure onset due to its downstream effects (as illustrated by a “GABAergic initiation hypothesis” schematized in Fig 1).

We also highlight an important distinction between this work and other computational work investigating the role of GABAergic signalling in epileptiform activity and inter-ictal discharges (IID): while recent literature investigating this topic makes use of the potential depolarizing capacity of GABA [56, 57], the work presented here uses purely inhibitory GABAergic synapses. Indeed, while changes in the GABA reversal potential are seen during seizure *propagation* [3], the changes in chloride concentrations necessary to elicit this feature are unlikely to exist prior to or during seizure *initiation*, which is the focus of this research. Moreover, the mechanisms proposed in the work of Chizhov et. al. [56, 57] do not focus on the capacity of excitatory cells for PIR, in contrast to the “GABAergic initiation hypothesis” discussed here.

The mechanism proposed in this paper is the first of which the authors are aware that describes how the initial step of a “GABAergic initiation hypothesis” might occur with both biological [12] and computational (this study) support. This, in turn, provides new and convincing evidence that may help to explain how the hyper-excitability induced by 4-AP causes the cortex to be more vulnerable to seizures, and more generally how interneurons can be involved in the initiation of cortical seizures clinically [58, 59].

### Mechanism details

The exploration of a transition driven by a small, permanent increase to the external drive was motivated by modeling studies [21] and physiological evidence [24] of inhibitory networks in the hippocampus. The observed sharp transition in the hippocampal model networks of Ferguson et. al. [21] was dependent on constraining the model network from cellular, synaptic and connectivity perspectives with the experimental data and context. The research presented here revealed that those hippocampal insights were translatable to a more generic, cortically-motivated network. Given this, it is possible that these insights are generalizable to most fast-firing inhibitory networks, although parameters representing external drive and synaptic strengths would not necessarily be the same. Additionally, considering the similarities in neural and network properties utilized in this work and that of Ferguson et. al. [21], it is very probable that the hippocampal networks would exhibit bistability of some form. However, of critical importance in the context of this study is the lack of an appreciable difference in the tendency for 4-AP and control model networks to exhibit this transition.

The mechanism articulated in this paper making use of the mathematical concept of bistability addresses the shortcomings, in the context of seizure initiation, of the mechanism described above. Bistability arises on a small scale in many neuron models, including the Hodgkin-Huxley equations, in which both the resting state and periodic firing of action potentials are stable solutions and the amplitude of the input to the system determines which of these dynamics is exhibited by the model [47]. Here, bistability was observed in the larger scale, network dynamics of coherent and incoherent network states. These states were uncovered by making use of a similar perturbation to that utilized previously in a more abstract study of inhibitory networks [23]. Critically, the transition from asynchrony to synchrony brought about by this mathematically-motivated perturbation persisted when a lower amplitute and wider perturbation, motivated by activity that might arise from an Ornstein-Uhlenbeck process simulating background excitatory synaptic activity, was used. This result indicates that this transition is potentially viable in a biologically-grounded setting as well.

The analysis of this “bistable transition” reveals that it is more likely to occur in 4-AP networks as opposed to their control counterparts. This result indicates that it is a much more likely culprit in the initial step of a “GABAergic initiation hypothesis” of seizure than a transition brought about by a small, permanent increase in external drive to the network. Further support for bistability serving a role in seizure onset is found in the statistics of inter-seizure intervals [60]. Moreover, the cortically-motivated Ornstein-Uhlenbeck process generated current displayed in Fig 6 illustrates that perturbation-like activity is more likely to arise from background synaptic excitation than longer-lasting increases approximating a permanent increase in the external drive to an inhibitory network. Taken together, these insights support the hypothesis that dynamical changes made possible by network bistability mechanistically explain how interneuronal populations are “hijacked” in pathology [61].

### The role of firing frequency

Given that the primary difference between the 4-AP and control model neurons is in the hyper-excitabillity of the 4-AP neurons, it bears investigating whether this feature plays a disproportionate role in dictating the overall network dynamics. To analyze this, a “Mean Firing Frequency” measure (which involves simply summing the total number of spikes in the network over a given time interval, dividing by the number of cells, and then converting this value into a frequency by dividing by the length of the time interval) was taken over the last 500 ms of simulations performed from random initial conditions over the parameter space used in Fig 4. The results, presented in Fig 9, reveal that the average cell firing frequency in control and 4-AP networks with similar network parameters and similar dynamical states (i.e. synchrony or asynchrony) are actually quite close (and any differences are certainly diminished from the extreme differences seen in their FI curves presented in Fig 2). This finding is fairly robust over all but the weakest inhibitory syanptic weights. Thus, the critical implication is that the mechanism involved in the “bistable transition” involves an interplay of cellular (potentially not only the hyper-excitability, but also the increased adaptation, in 4-AP neurons) and network properties, and could not be replicated merely by causing the neurons to fire faster in some artificial fashion.

**Fig 9.**
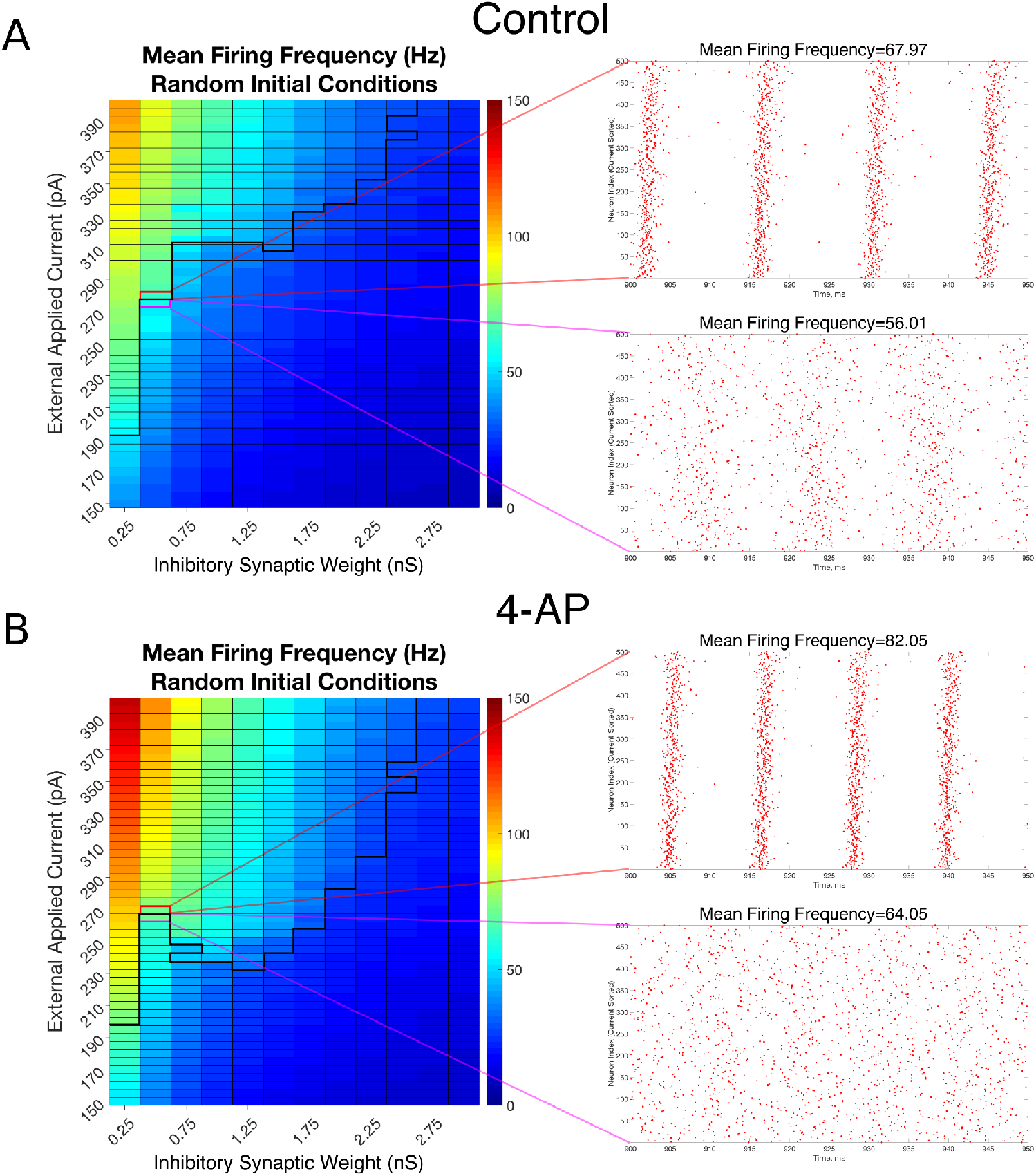
Overall firing frequency is higher in networks exhibiting synchrony, providing a potential avenue through which to identify the results of this work via experimental recordings. A-B: Mean Firing Frequency values, averaged over five independent simulations, for control (A) and 4-AP (B) networks. The border dividing the parameter regime supporting synchronous dynamics (top-left) from the regime of asynchrony is depicted by the bolded black line, where this border was found using a cutoff value for the Synchrony Measure of 0.25 (which was found to be reasonable after a rigorous investigation of a variety of raster plots and their corresponding Synchrony Measures). Example raster plots for the networks outlined in pink and red illustrate example asynchronous and synchronous raster plots, respectively, along this border. Their Mean Firing Frequency values illustrate the relatively large increase in network activity that is associated with the transition from asynchronous to synchronous firing.

Also of interest in this analysis was that both control and 4-AP networks show a similar increase in average firing rate when transitioning from asynchrony to synchrony (highlighted by the example raster plots and corresponding mean firing frequencies presented in Fig 9). This result is analogous to a similar finding in [21] and indicates a potential avenue for an experimental exploration of the results presented here: namely, a substantial increase in firing rate following a perturbation to an inhibitory network is likely indicative of the transition into synchronous firing. Such behavior is likely more easily identifiable by multi-electrode arrays than synchronous firing itself.

Furthermore, these results shed new light on the interaction between synchrony and increased cell firing rate in the context of seizure onset. Experimental literature commonly shows that these dynamics (in both excitatory and inhibitory cells) both accompany seizure onset (see, for example, the work in humans of [62]), with many of these studies implying that increased firing rate plays a causal role in the transition into synchrony (see, for example, the work of [13] which reveals an increase in interneuron firing rates prior to seizure and the corresponding synchronous dynamics). However, the “bistable transition” described in this paper does not require a change to the system that would increase the average cell firing rate; rather, the increased firing rate comes about seemingly driven by the induced synchronous firing of the inhibitory network. Thus, it is possible that synchrony of inhibitory networks is permissive of an increased neural firing rate, instead of increased firing rate causing this synchrony. Indeed, where there is sparse sampling of interneurons, increased firing rates of interneurons prior to a seizure may be additionally interpreted from our modelling results to represent a transition to synchronous interneuronal firing [13] rather than a firing rate increase alone.

### Theoretical insights and related studies

The multi-scale and nonlinear nature of the human brain makes it challenging to understand its dynamics. As such, insights from theory are needed to help guide computational studies and inform the understanding of brain networks. Here, models of inhibitory networks informed by cortical data were used to explore potential mechanisms leading to the transition from asynchrony to synchrony that occurred more robustly in hyper-excitable settings. Such synchrony primarily corresponded with fast network oscillations.

However, networks of fast-firing interneurons can also produce slow population output as shown in modeling studies [63]. The ability of fast-firing inhibitory networks to produce slow population activities was shown to be possible via individual cells having enough of a “kink” in their frequency-current (FI) curves that allowed a bistable network mechanism to be present [63]. The modeled slow population activity (< 5 Hz) is seen *in vitro* using a hippocampal preparation [64, 65], and a bistable network mechanism was subsequently leveraged to explain paradoxical changes seen in Rett syndrome mice from the perspective of these same slow population activities [66].

A critical difference between the bistable network mechanism of Ho et. al. [63] and bistability related to properties of the ING mechanism (analogous to that presented here) was summarized by Skinner and Chatzikalymniou [67]. In the work of Ho et. al. [63], the mean excitatory drive received by inhibitory cells in the network must be close to their spiking rheobase. The bistability is between states with low or high numbers of fast-firing cells, and this allows slow population activities to come about due to excitatory fluctuations in the system. A similar mechanism could be in play in the work of Schlingloff et. al. [24] where an *in vitro* representation of sharp waves was examined and it was suggested that sharp waves could be generated stochastically from excitatory input. In contrast, for an ING-related bistability, the excitatory drive to the inhibitory cells is not close to spiking rheobase, but as shown by Rich et. al. [23] and in the cortically-motivated networks presented here, bistability between synchronized high frequency firing and asynchrony is possible.

There have been numerous studies in the computational literature probing the tendency for networks of inhibitory neurons to synchronize, although these studies typically are done in a more theoretical setting rather than the biologically-motivated manner presented in this study. The interneuron models utilized here exhibit Type I properties in their FI curves (namely, a steep FI curve with an arbitrarily low firing frequency [68]), and neurons with these properties have been a focus of many computational studies of inhibitory synchrony [69–72]. As such, the coherent dynamics seen in our cortically-motivated inhibitory networks correspond with insights from these more abstract computational studies. This literature contributed to the articulation of the ING mechanism [15, 16, 18, 19] that is most likely driving the coherent dynamics seen in these cortical inhibitory networks. Another seminal study on inhibitory synchrony and ING found that the synchrony promoted by the ING mechanism is most robust when networks are more densely connected and cellular heterogeneity is low [73], features replicated in the cortically-motivated networks presented here.

In this context we note that computational studies proposing mechanisms for synchronous network oscillations are typically concerned either with purely inhibitory networks (as presented here), purely excitatory networks [74], or networks containing inter- and intra-connected subnetworks of excitatory and inhibitory cells (E-I networks). Crucially, the mechanisms underlying synchrony and their dependence on features such as cell excitability properties (i.e. the Type I vs. Type II distinction [68]), external drive to the network, and network connectivity can vary significantly depending on the type of network studied. For example, the results of Hansel et. al. [74] imply that an excitatory network made up of cells of the type studied here is highly unlikely to ever synchronize. Similarly, while the Pyramidal Interneuron Network Gamma (PING) mechanism is commonly cited as a mechanism causing synchronous oscillations in E-I networks [16, 72, 75, 76], recent work has revealed that the predictions of this mechanism are altered by varying individual cellular properties and network connectivity [42, 43].

### Limitations and Future Work

The neuron models implemented here used a simplified Izhikevich type integrate and fire mathematical structure, informed by a combination of existing literature and in-house experiments. With a full repertoire of experimental recordings, one could more fully capture neuronal features and differences, but a consideration of the multiple inhibitory cell types as well as network configurations and properties should also be taken into consideration. Indeed, one could consider designing a neuromodulation study using the Blue Brain Project [37] to examine this given the insights gleaned from this study.

The network structure used in this work, a purely inhibitory network, is also simplified from the biology. However, this choice was critical in allowing for the articulation of a mechanism of action underlying the transition of interneurons into synchrony. Such a mechanism is a paramount and necessary “first step” towards an overarching mechanism of a “GABAergic initiation hypothesis” and provides initial justification for further, more biologically detailed study of this hypothesis. With this mechanism in hand, future work can more easily investigate how the dynamics of excitatory cells might affect or interact with this behavior amongst the inhibitory neurons.

For the work here, we focused on differences between control and 4-AP neurons as encapsulated in our models. It is unlikely that utilizing a more realistic noisy synaptic input would affect the primary results of this work, since both noisy [77] and deterministic [21] inputs were used in previous hippocampal inhibitory network models without changing insights regarding the transition into synchrony.

While use of a simplified neuron model and network structure enables extensive parameter explorations to be easily done and dynamical aspects, like bistability, to be uncovered, parameter interpretation relative to details of the biological system is less straightforward. However, studies such as this could help leverage understanding and motivate hypothesis-driven explorations in more detailed models. We note that due to the relatively sparse connectivity of the cortically-motivated inhibitory networks studied, mathematical tools such as reduction to phase oscillator models as in Hansel et. al. [74] that require an assumption of all-to-all connectivity and weak coupling cannot be easily applied to do further theoretical analyses.

With the ability to obtain and model human cellular data [78, 79], it will be interesting to consider how one might examine and take advantage of these mechanistic insights. Expanding our mechanistic insights here to include excitatory cell networks and other seizure phases (see Fig 1 [5]) are exciting considerations.

## Conclusion

The hypothesis that excessive inhibitory signalling serves a causal role in seizure initiation has support from numerous recent experimental studies, but until now has not subject to rigorous computational investigation. Here, we provide the first computational support for this theory by articulating a mechanism that not only potentially explains the necessary, initial step in this process (the sudden transition of inhibitory interneurons from asynchronous to synchronous firing), but also why networks in a model ictogenic state are more vulnerable to this transition. This latter feature distinguishes this mechanism as one particularly important in the study of epilepsy, as any process asserted to relate to seizure initiation should be more likely to occur in an ictogenic as opposed to healthy brain. As this novel hypothesis of seizure initiation is inherently counter-intuitive, especially in comparison to the more historically common theory of over-excitation or dis-inhibition initiating seizure, this computational work articulating potential mechanisms of action is especially important to support the theory’s viability. Indeed, our work presented here not only provides the first evidence of which we are aware that feasible mechanisms underlying necessary steps in this process exist, providing paramount support for a “GABAergic initiation hypothesis”, but also presents new potential avenues for experimental and clinical epilepsy research via further investigation of this mechanism.

## Acknowledgments

We thank Amir Rez Peimani for early simulations performed for this work. Funding for this work was provided by the Natural Sciences and Engineering Research Council of Canada (NSERC) via Discovery Grants to Frances K. Skinner and Taufik A. Valiante.

